# The folding-limited nucleation of curli hints at an evolved safety mechanism for functional amyloid production

**DOI:** 10.1101/2023.05.26.542396

**Authors:** Jolyon K. Claridge, Chloe Martens, Brajabandhu Pradhan, Frank Sobott, Mike Sleutel, Han Remaut

## Abstract

It is nearly two decades ago that the ‘thin aggregative fimbriae’ which had been shown to enhance the biofilm formation of *Salmonella enteriditis* and *Escherichia coli* were identified as amyloid fibers. The realization that natural proteins can develop amyloidogenic traits as part of their functional repertoire instigated a search for similar proteins across all kingdoms of life. That pursuit has since unearthed dozens of candidates which now constitute the family of proteins referred to as functional amyloids (FA). FAs are promising candidates for future synthetic biology applications in that they marry the structural benefits of the amyloid fold (self-assembly and stability) while steering clear of the cytotoxicity issues that are typically linked to amyloid associated human pathologies. Unfortunately, the extreme aggregation propensity of FAs and the associated operational difficulties are restricting their adoption in real-world applications, underscoring the need for additional processes to control the amyloid reaction. Here we untangle the molecular mechanism of amyloid formation of the canonical functional amyloid curli using NMR, native mass spectrometry and cryo-electron microscopy. Our results are consistent with folding-limited one-step amyloid nucleation that has emerged as an evolutionary balance between efficient extracellular polymerization, while steering clear of pre-emptive nucleation in the periplasm. Sequence analysis of the amyloid curlin kernel suggests a finetuning of the rate of monomer folding via modulation of the secondary structure propensity of the pre-amyloid species, opening new potential avenues towards control of the amyloid reaction.

## Introduction

Amyloids encompass a unique class of filamentous structures that emerge from a broad range of self-assembly routes. Once characterized by a single defining structural feature, i.e. the cross-beta architecture^1^, it is now clear that the structural diversity of amyloid filaments is matched by the variety of primary protein sequences that can adopt said amyloid fold^2^. For a long time, interest in amyloids has been driven by their connection to a host of (neuro)degenerative misfolding and deposition diseases^1^, and more recently, by their function in a range of biological processes such as adhesion, storage, biofilm scaffolding, etc^3, 4^. The dichotomy between so called “pathological” and “functional” amyloids, or also “incidental” and “evolved” amyloids, exposes a blind spot in our understanding of amyloids and raises questions regarding the precise origins of their reported cytotoxicity, and the corresponding mitigation strategies of the cell. Progress on these fronts has been made by deciphering the succession of molecular events that take place between the monomeric starting point and the emergence of the fibrous amyloid ultrastructure, and any off-pathway routes that may intersect^4–6, 7^. Most pathological amyloids represent a non-native folding state that originates from destabilized or alternatively spliced or fragmented proteins and peptides. The amyloid assembly route encompasses the rapid formation of reversible, non-native oligomers (pre-nuclei) and aggregates that undergo a rate-limiting structural reconversion into an amyloid protofibril nucleus that serves as a templating catalyst of a non-native polymerization process. Amyloid toxicity can sometimes be ascribed to membrane destabilizing, thinning and/or pore forming activity of such dynamic pre-nuclei species^7^. There are clear indications that mature amyloid fibers can affect organ integrity, by instigating inflammatory responses or by membrane disruption and lipid extraction by fibril termini^8^. What molecular and structural aspects of amyloid assembly pathways or fibril structure determine cytotoxicity and disease often remain poorly understood. Case in point are recent structural studies of tau filaments extracted from patients with different tauopathies, and which were found to be associated with unique tau filament folds^9^. How such filament polymorphs or their different assembly pathways relate to disease outcome is unclear.

Here we turn to study bacterial functional amyloids as model systems of amyloid deposition pathways evolved to minimize cytotoxic activity towards the host cells. For obligate functional amyloids, where the amyloid state makes up the sole native state of the protein^3^, one can expect positive selection to shape the nucleation and polymerization pathway(s) and negative or purifying selection to minimize the cytotoxicity of such processes. Arguably the best studied case is that of curli. Curli fibers are a major component of the extracellular matrix of Gram-negative bacteria and are expressed under biofilm forming conditions where they fulfill a scaffolding and cementing role in the extracellular milieu to reinforce the bacterial community^10^. The major subunit of *Escherichia coli* curli is the 13.1 kDa pseudo-repeat protein CsgA, which is secreted as a disordered monomer that folds into β-solenoid domains that stack head-to-tail to produce non-helical, polar, protofilaments^11^ with classic amyloid tinctorial properties, under a wide range of conditions^12, 13^. In its native context, formation of cell-associated CsgA fibrils requires the minor curli subunit CsgB, which in turn is bound to the outer membrane CsgG-CsgF secretion pore complex^12, 14^. Although CsgB also acts as a nucleation seed *in vitro*^15^, purified CsgA monomers will readily form native-like amyloid curli fibers^16^. This intrinsic CsgA nucleation occurs within minutes and at low nanomolar concentrations^16^. Global fitting analyses of Thioflavin T (ThT)-fluorescence kinetics, showed that *in vitro* curli formation can be modelled with a simple model for homogenous nucleation coupled to elongation, ruling out self-replication as an important source of nucleation^17^. Based on the time-dependent reactivity of CsgA aliquots with the Ab-oligomer specific antibody A11, Wang et al^18^ argued the existence of transient folding intermediate CsgA species, suggestive of a curli polymerization mechanism that shares similarities to the nucleated conformational conversion process (NCC) pathway of pathological amyloids^5^. However, using *in situ* imaging at the nanoscopic level, we previously excluded the involvement of aggregated or oligomeric pre-nucleus assemblies akin to those observed for the NCC mechanism of pathological amyloids^16^. This study found a direct nucleation pathway forming minimal amyloid species that showed elongation kinetics equivalent to those of mature fibrils within the seconds time-resolution of the AFM imaging technique.

What remains to complete our picture of *de novo* curli formation is an identification of the nucleus species. In a recent NMR study of pre-amyloid CsgA, Sewell *et al* did not find any signs of any folded sub-populations, suggesting that such species are either rare, short-lived, or both^19^. Although ensemble techniques such as NMR and ThT fluorescence can provide insights into the structural details of the pre-amyloid state and kinetic properties of amyloid conversion, respectively, they are not well-suited to probe minority or individual events. Here, we adopt an approach catered to the stochastic nature of nucleation by combining two techniques that can kinetically trap and probe the rare species that emerge in the early stages of curli formation. Based on native mass spectrometry (nMS) analysis and cryogenic electron microscopy (cryoEM) imaging of polymerizing CsgA solutions, we identify the smallest folded CsgA species to be a β-solenoid dimer. This suggests that the curli-formation is not a nucleation-limited process in the classical sense of producing a metastable nucleus of critical size. Rather, sigmoidal ThT-kinetics point towards an initial, slow folding step that places constraints on the rate of fibril formation. Taken altogether, a picture emerges of folding-limited one-step nucleation that has emerged as an evolutionary balance between efficient extracellular polymerization, while steering clear of pre-emptive nucleation in the periplasm.

## Results

### CsgA exhibits bi-phasic switching from unfolded monomers to folded amyloid fibrils *in vitro*

Using dynamic light scattering, ThT-fluorescence, and circular dichroism spectroscopy experiments on nucleating CsgA solutions, we and others have previously shown that CsgA fibril formation follows a monotonic, sigmoidal kinetic dependence^16, 20^. These biophysical methods probe the fraction of CsgA that has accumulated in the insoluble, amyloid state. To gain insight into the soluble, pre-amyloid state, we employ solution-state NMR on samples that contain unfolded CsgA monomers at the onset. For this we start from recombinantly produced CsgA that was purified from inclusion bodies under denaturing conditions (8M urea). We allow spontaneous amyloid formation to set off by exchanging the chaotropic storage buffer (25mM K-Phosphate 7.5, 8M Urea, 250 imidazole, 50mM NaCl) with 15mM MES pH 6.0 via passage over a Hitrap desalting column immediately prior to NMR data collection. Next, we recorded 1D proton spectra at one-minute intervals over a time-course of 12 h. These time-resolved spectra exhibit no chemical shift changes but gradually decrease in intensity over time (Figure 1a). Next, we integrated the signal over a 0.5 ppm window in the 8-8.5 ppm amide region and plotted the intensity as a function of time (Fig.1a inset). The resulting signal is a proxy of the soluble CsgA fraction and exhibits a monotonic temporal dependence, reciprocal to the appearance of CsgA fibers as measured by ThT fluorescence (Supporting Figure 1). This fits with a model of fibrillation involving a simple two-state conversion process, going from a soluble, monomeric pre-amyloid disordered state to a folded fiber-incorporated amyloid state. Any intermediate or alternate conformations would be short-lived and/or sparsely populated. To widen the temporal window on the earliest stages of nucleation, we collected a similar kinetic trace for a four-point mutant^18^ of CsgA^Q49A/N54A/Q139A/N144A^ (hereafter referred to as CsgA^slowgo^ ; Fig.1a inset) which is characterized by slower fibrillation kinetics but produces otherwise identical curli amyloid fibers^21^. Our data show that the CsgA^slowgo^ signal also decreases monotonically as a function of time, albeit at a slower rate in line with earlier ThT-data^21^, resulting in little loss of monomer signal in the initial 40h. To gain further molecular insights into the solution state during this lag-period, we collected and assigned HSQC spectra for CsgA^slowgo^. A representative HSQC spectrum is shown in Fig.1b and is very narrow in the proton dimension (0.7 ppm), indicative of IDP-like characteristics of the CsgA^slowgo^ molecules. The furthest downfield shifted backbone amide proton resonance is that of N42 at 8.51 ppm, with the furthest upfield-shifted backbone amide proton (except for the C-terminal Y151 at 7.76ppm) of I47 at 7.81 ppm. For comparison, the predicted backbone amide shifts of a β-solenoid conformation fall between 7.01 and 10.55 ppm (Supporting Figure 2).

**Figure 1:**
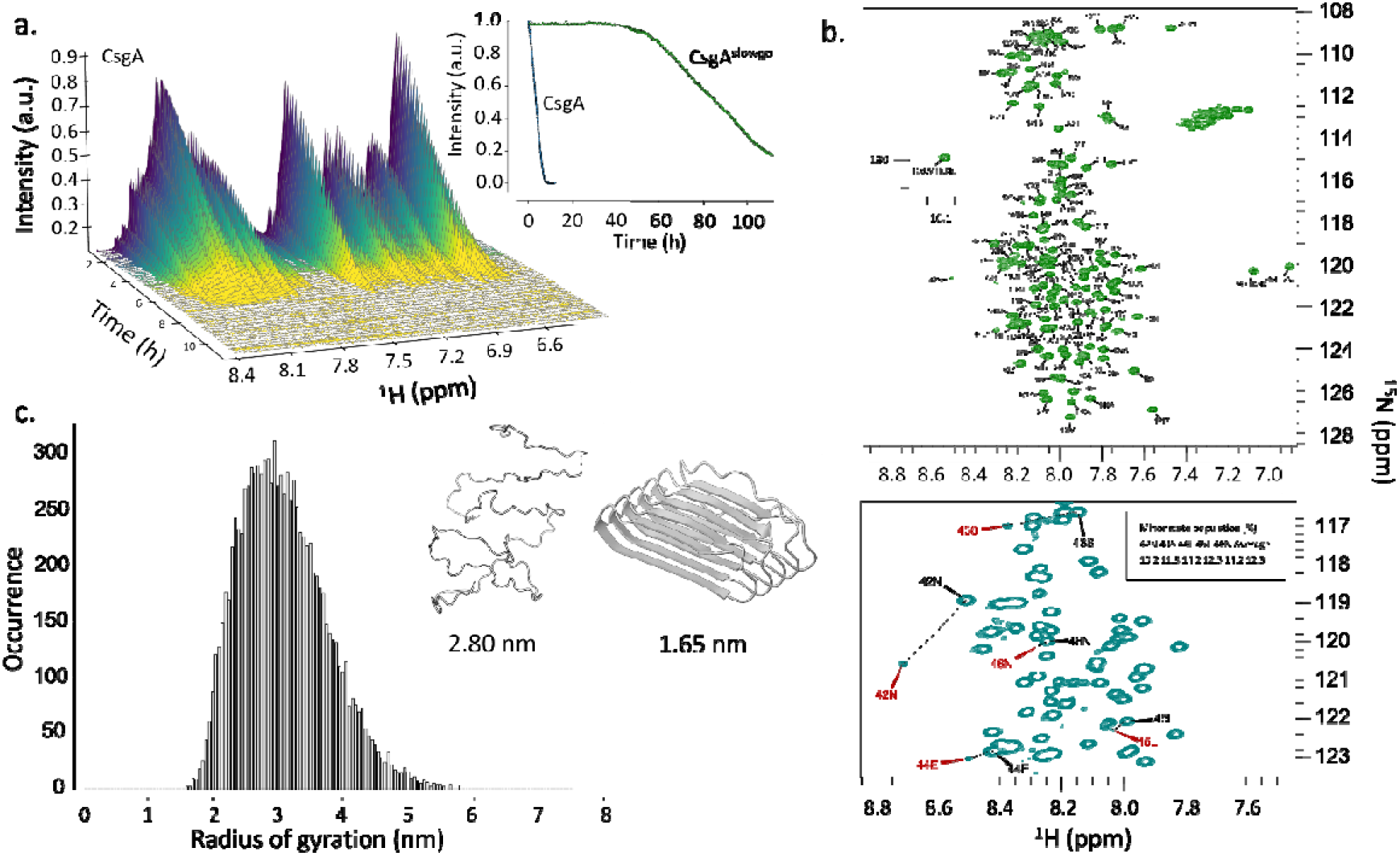
**(a)** Waterfall plot of NMR signal intensity over time for CsgA (main figure) and for an integral of the amide region of a 1D proton spectrum of CsgA and CsgA^slowgo^ over time (inset). It can be seen that the signal for CsgA has decayed to zero after approximately 8 hours, whereas the signal is still observable for CsgA^slowgo^ after over 100 hours in the spectrometer. (**b**) ^15^N-HSQC spectrum of CsgA^slowgo^ showing assigned residues. The inset shows the HeNe from W106 which is downfield shifted outside the displayed spectral width. The second panel shows a region of ^15^N-HSQC spectrum of CsgA^slowgo^ showing peaks that exhibit doubling (between N42 and N46), indicating a minor population probably due to proline isomerisation. (**c**) Distribution plot of radii of gyration for members of a Flexible Meccano generated ensemble of CsgA (main panel) and representative example of the ensemble rendered in PyMol with the average radius of gyration and an AF2 model of the putative folded conformation of CsgA with its radius of gyration.

To gauge the hydrodynamic radius (*R_h_*) of CsgA^slowgo^ in this pre-amyloid state, we performed DOSY experiments from which we obtain a diffusivity of 1.37 x 10^-10^ m^2^s^-1^. Using the Stokes-Einstein equation, we calculate *R_h_* to be 1.82±0.03 nm which is in line with what can be expected for a molecule of 13 kDa. To put this in perspective, we estimated the distribution of the radius of gyration (Fig.1c) derived from an ensemble of unfolded CsgA polypeptides generated using Flexible Meccano^22^. The mean *R_g_^unf^* of this distribution is 2.8±0.8 nm, i.e. significantly larger than the experimental *R_h_* estimate. Conversely, we calculated *R_g_^fold^* of a folded CsgA monomer using an AF2 predicted β-solenoid monomer as a template^11^. The predicted *R_g_^fold^* for this species, 1.65 nm was calculated by HYDRONMR^23^ which is relatively close to the experimental estimate derived from DOSY. Next, we gauged the foldedness of monomeric CsgA^slowgo^ by determining the water – amide proton exchange rates using the CLEANEX approach, and comparing these with predicted rates for CsgA^slowgo^ in random coil conformation, as calculated by the SPHERE server (https://protocol.fccc.edu/research/labs/roder/sphere/sphere.html24). The highest amide exchange rates were measured for N22 (Fig. 2a), a 22-residue sequence in mature CsgA that acts as a targeting sequence for CsgG-mediated secretion^25^, but is absent from the curli amyloid core^12, 13^. For the most part, amide exchange rates in N22 residues approach those predicted for a random coil, indicating that this region is largely unstructured. In contrast, backbone amide exchange rates in the curlin repeats significantly deviated from calculated random coil values (Fig. 2a). Experimental rates averaged to 0.69 s^-1^, close to the detection limit of 0.5 s^-1^, compared to an average 3.0 s^-1^ for the random coil. The highest rate predicted by SPHERE is 16.5 s^-1^ for residue G74 in R2, the corresponding experimental value is 4.5 s^-1^. This increased shielding of the backbone amides from exchange with the bulk solvent suggests the presence of at least transient backbone-backbone and/or backbone-sidechain interactions in the curlin repeats.

**Figure 2:**
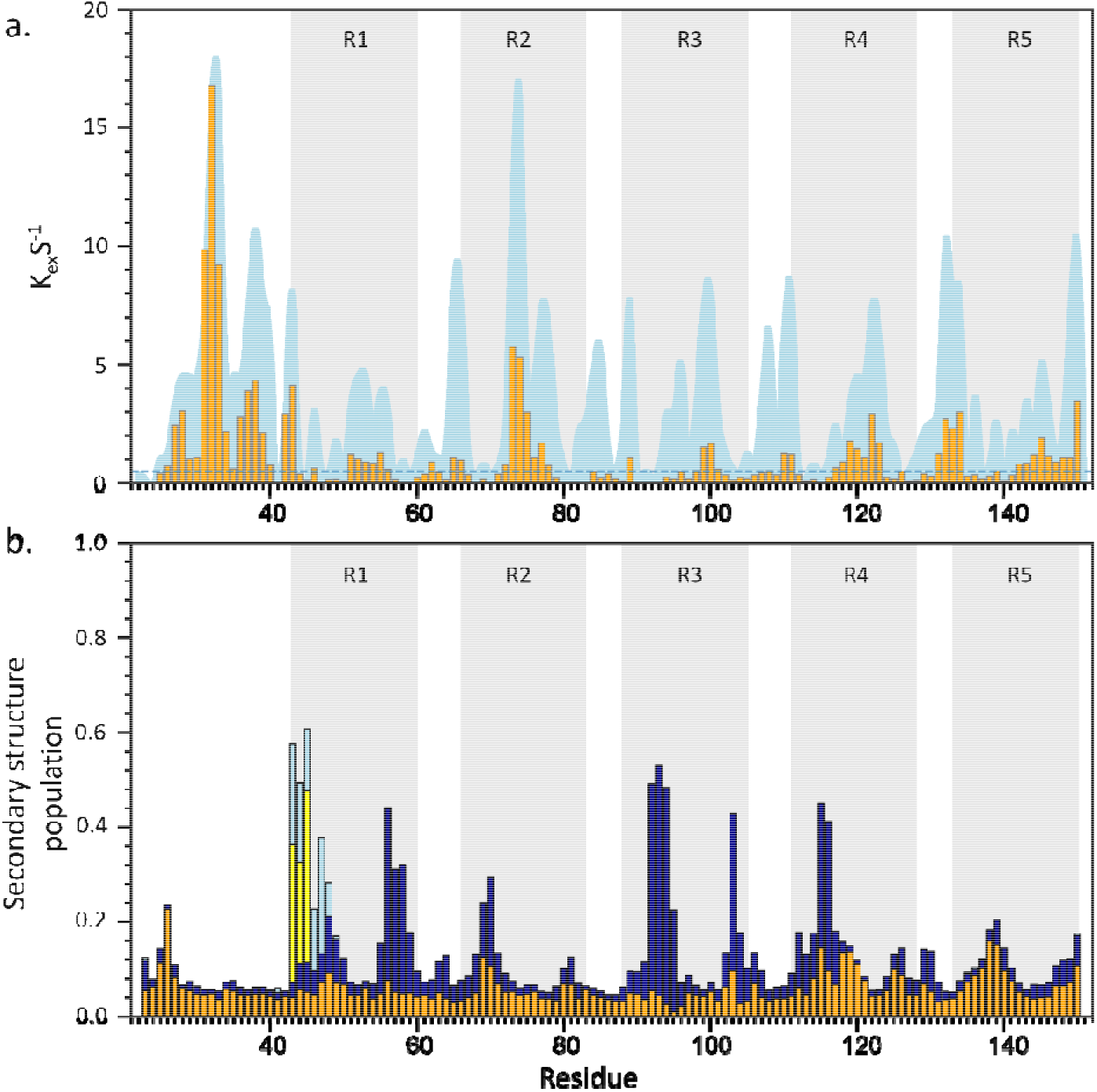
**(a)** CLEANEX^50^ derived amide exchange rates for CsgA^slowgo^ plotted for each residue (orange bars) with the SPHERE predicted rates for CsgA^slowgo^ (blue shading). The curlin amyloid repeats are shaded grey. The minimum detectable exchange rate of 0.5 s^-1^ is depicted as a blue dotted line. **(b)** TALOS derived secondary structure propensity for CsgA^slowgo^ major and minor populations. Blue bars represent helical propensity (either α-helix or PPII helix) and orange represents β-strands. The differences between the two populations are between P41 and N46, these are shown as light blue and yellow bars for the helical and β-strand propensities respectively.

To obtain further structural insights into the pre-amyloid state of CsgA^slowgo^ we used TALOS-N to predict backbone and sidechain torsion angles (Fig. 2b). Remarkably, thusly derived secondary structure propensities of CsgA^slowgo^ reveal a weak (α-or PPII-) helical character of the sequence motifs that give rise to the strand-turn-strand architecture of folded CsgA monomers inside the curli fibers^11^. More specifically, in the pre-amyloid monomers, repeats 1 to 5 all exhibit helical propensity, though there is some tendency for β-strand in the C-terminal repeats (R4 and R5). Moreover, the minor population related to the cis/trans isomerisation of Pro41 shows a strong but local strand tendency between residues N42 and N46. The chemical shifts of P41 Cα and Cβ in the minor population are shifted downfield (0.66 ppm) and upfield (2.57 ppm) respectively (Supporting Fig.3). This indicates *cis* conformation of the proline in this population. Taken altogether, these results suggest that CsgA^slowgo^ is neither folded nor fully extended, but rather adopts a molten-globule like state, sampling local, mostly helical secondary structure.

**Figure 3:**
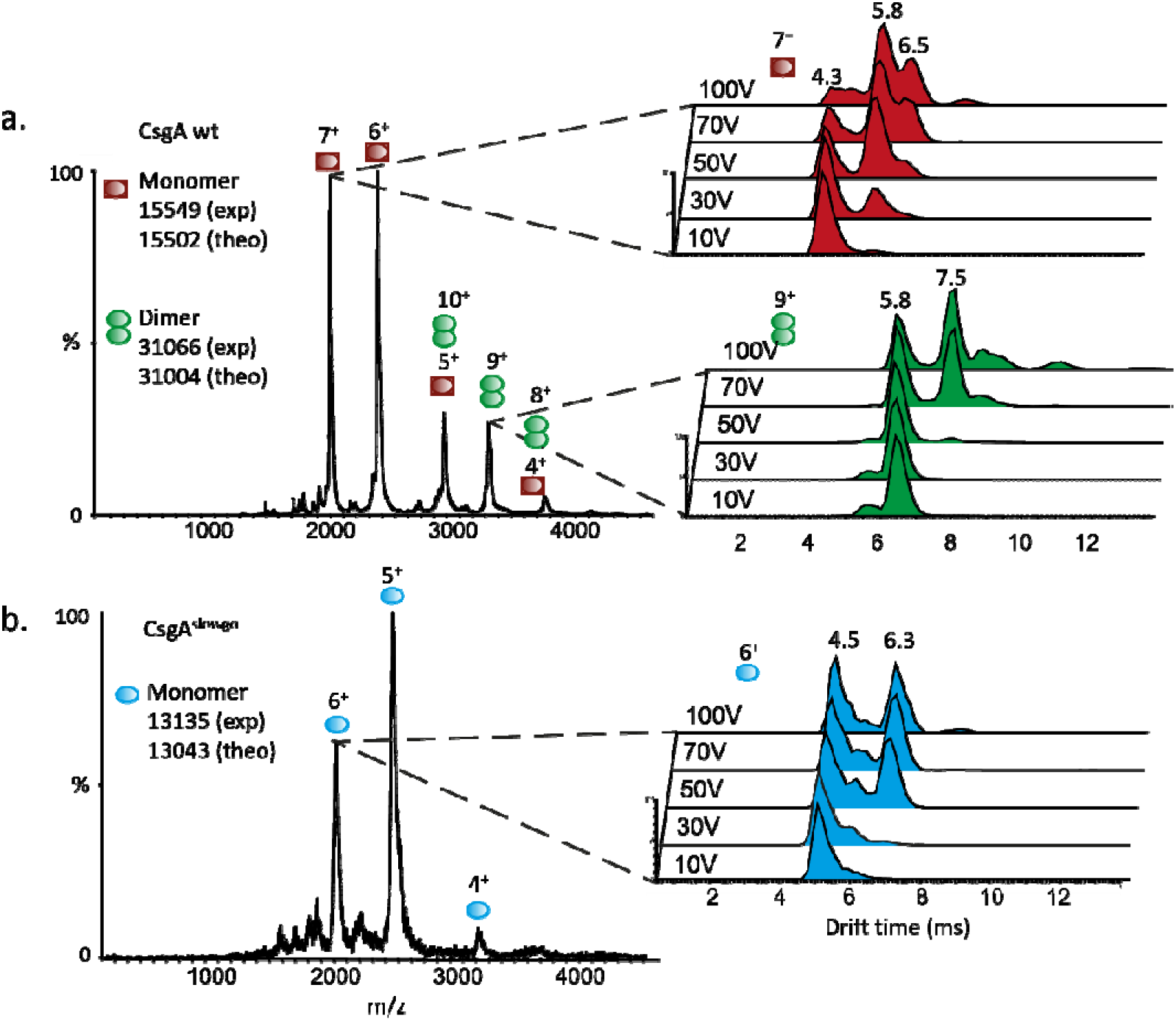
The ESI mass spectra of CsgA wt **(a)** and CsgA^slowgo^ **(b)** reveal the presence of monomeric (red) and dimeric CsgA wt (green) species while CsgA^slowgo^ is observed only as a monomer (blue). The insets show the drift time of monomeric CsgA wt (red), dimeric CsgA wt (green) and monomeric CsgA^slowgo^ (blue).

### CsgA forms dimers during the early stages of fibrillation

We did not identify a clear signature of the nucleating species in our NMR data, which is likely related to (1) the paucity, (2) the short-lived nature, and/or (3) excessive size of said species. To increase our sensitivity towards rare and short-lived nucleation events, we adopted a non-ensemble methodology that can accurately resolve various condensation and oligomeric states of CsgA under near-native conditions. For this we used native ion mobility mass spectrometry (nMS) to follow the oligomerization dynamics of CsgA as a function of time. Both wild-type (WT) and CsgA^slowgo^ were buffer exchanged from 8M urea to a nMS compatible buffer (150mM ammonium acetate pH7) immediately before analysis by nano-electrospray ionization coupled to ion-mobility mass spectrometry (nESI-IM-MS).

Monomeric CsgA was readily observed both for the WT sample and CsgA^slowgo^ (Fig 3 and Supporting Figure 4). In the WT sample, a second species was observed corresponding to dimeric CsgA (Fig 3a). We performed collision induced unfolding experiments to evaluate the relative stability of the different species observed (expanded peaks in Figure 3). We observed an “unfolding”-like transition for all three species, characterized by a shift in the drift time distribution from shorter to longer times. From this we hypothesize that the compact state is the native state of CsgA from solution, and the extended state is produced by gas-phase unfolding. Clear differences are observed between the monomeric forms of WT compared to CsgA^slowgo^. In the case of the WT, the compact state decreases substantially upon increasing collision voltage energy, while in the case of CsgA^slowgo^, significant levels of the compact state persists at higher collision energies. Similarly, the dimeric form of WT CsgA presents a compact state that is stable in the gas-phase even at collision energies as high as 70 and 100V, indicating that the dimeric form of WT CsgA is more stable than the monomeric form. It is worth noting that leaving the WT sample in native MS buffer on ice for 1 hour compromised spraying. In contrast, the CsgA^slowgo^ sample could still be sprayed after 2h on ice and the spectra obtained right after buffer exchange were very similar to the those obtained after 2h (Supporting Figure 5). These differences likely reflect the variation in oligomerization kinetics: once oligomerization has progressed too far, insoluble aggregates are formed and the sample can’t be efficiently ionized and transferred into the gas-phase.

**Figure 4:**
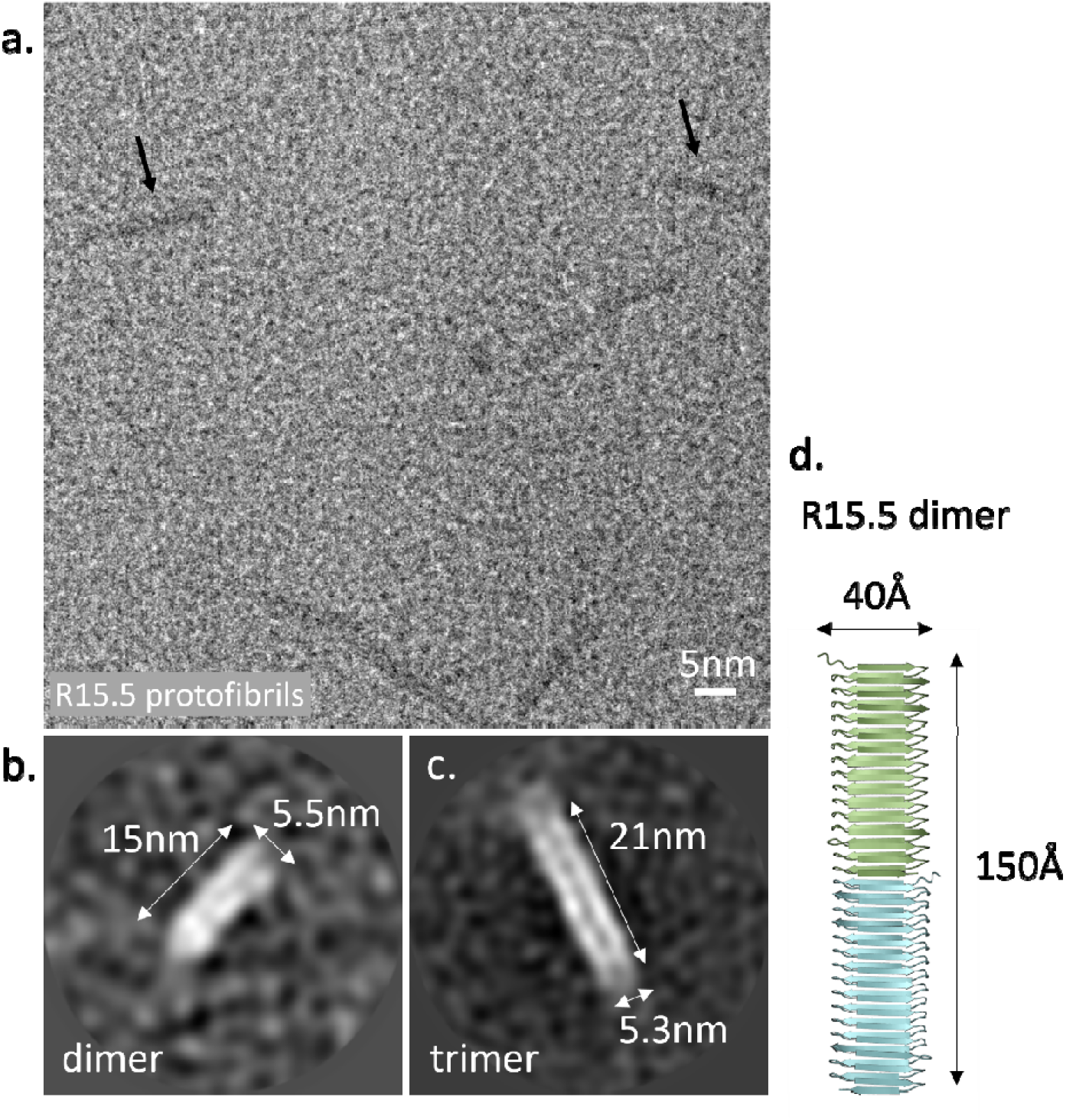
**(a)** Raw cryoEM micrograph of R15.5 protofibrils (black arrows as example) formed 2.5 min after desalting, deposited on a 2nm continuous carbon grid, dry-blotted and plunge-frozen in liquid ethane. **(b)** and **(c)** Relion 2D class averages of 1572 manually picked protofibril particles, tentatively assigned as R15.5 dimeric and trimeric species based on the indicated dimensions and reference^31^. **(d)** Cartoon representation of an AlphaFold2 model of an R15.5 dimer.

**Figure 5:**
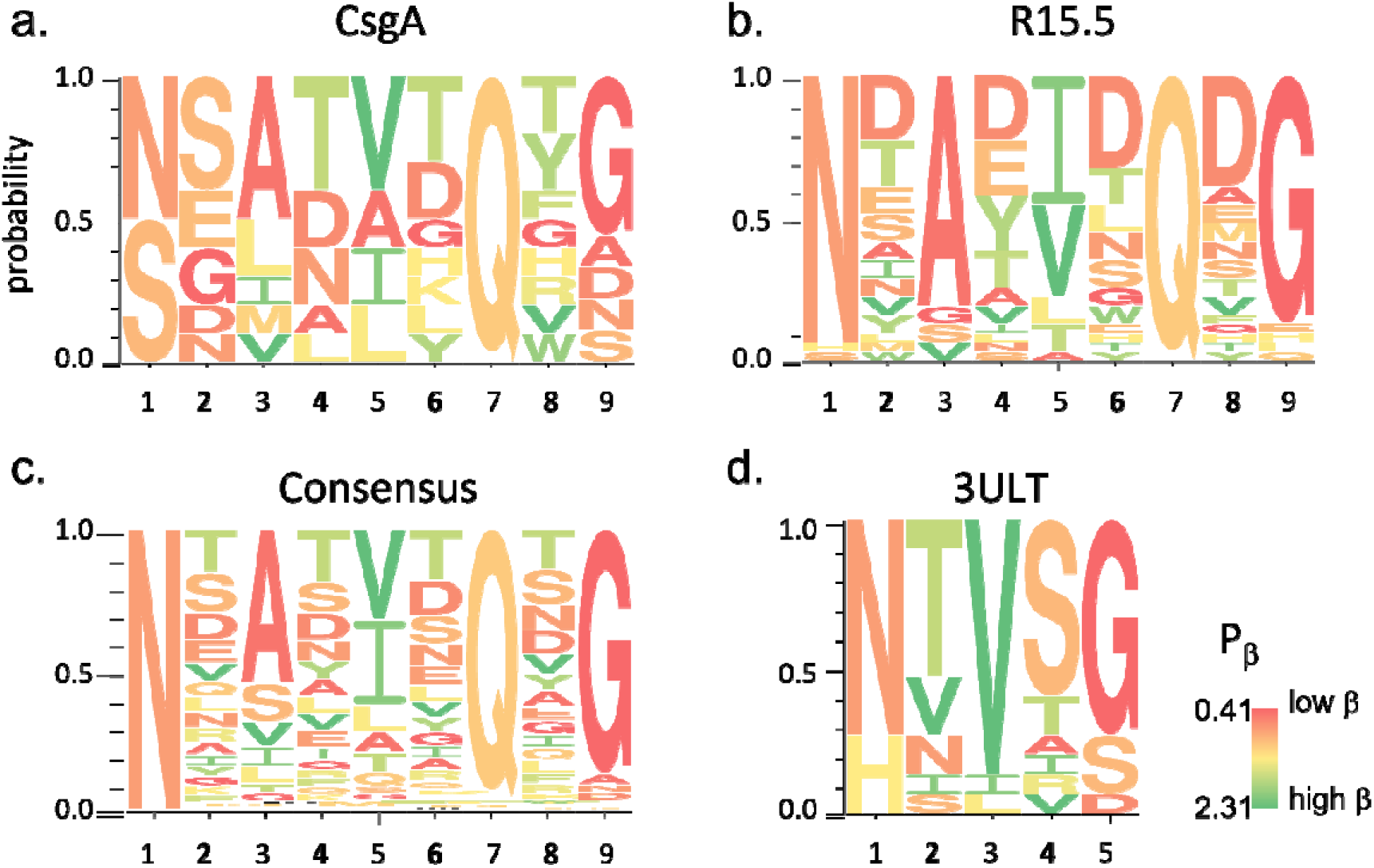
Amino acid frequencies for CsgA **(a)** and R15.5 **(b)** found along the 8 positions of a curlin strand, averaged across all repeats. **(c)** The consensus panel was constructed from a multiple sequence alignment of 53000+ repeat sequences that were mined from the refseq database. **(d)** 3ULT is a β-solenoidal ice-binding-protein that serves as a non-amyloidogenic reference. The colour coding in panels (a) to (d) is according to the average β-strand conformation propensities P_β_ taken from^32^ – green and red corresponds to high and low β-strand propensity, respectively.

Interestingly, we find that the dimer peak intensity reaches a maximum around 40min post desalting (Supporting Figure 6). After 60 minutes, CsgA monomers or dimers can no longer be detected. This temporal dependence supports an interpretation that the dimers are on pathway and not irreversible, dead-end artefacts of the transferal to the nESI gas phase. Similarly, it also demonstrates that dimers are not products of dissociation from larger oligomers at the selected trap voltage. If that were the case, one would expect the dimer peak to gradually accumulate as a function of time mirroring the amyloid number density in the liquid. Recently, a similar time-dependent dimer signal was also observed by Bhoite *et al*^26^. They recorded significant differences in the rate of attenuation of the peaks corresponding to the dimer species for CsgA homologs with different rates of aggregations.

### CryoEM resolves minimalistic amyloid nuclei composed of two or three CsgA molecules

Our previous high speed AFM imaging observed a direct nucleation pathway, corresponding to diminutive, short-lived species that adopted mature fiber elongation kinetics within the time-resolution of the measurements^16^. It is tempting to speculate at this point that dimeric CsgA species observed in nESI-IM-MS correspond to these early amyloid nuclei that catalyze fibril growth. Although the observation of a maximum in dimer density early in the polymerization experiments could be in agreement with this notion, we sought to gain further insights into the structural details of the early onset CsgA dimers using cryoEM. CryoEM is well-suited for this because it is not hampered by the short-lived nature of the reacting entities and can cope with the intrinsic stochasticity of nucleation^27^.

As *E. coli* CsgA dimers (4x2x3.5nm^3^) are currently below the size limit for cryoEM, we worked with a CsgA homologue that is composed of 15.5 curlin repeats, for which the predicted dimer (14x2x3.5nm^3^) is expected to produce sufficient contrast for particle picking in raw micrographs. We have recently shown that R15.5 self-assembles into curli protofibrils that are architecturally equivalent to CsgA fibrils, but are less prone to lateral aggregation and therefore more amenable to single-particle cryoEM^11^. Similar to our workflow for CsgA, we maintain soluble stocks of unfolded R15.5 monomers in 8M urea and allow for polymerization to initiate via a rapid desalting step to 15mM MES pH 6.0 (Supporting Figure 7). Two minutes after desalting, we apply a 3µl aliquot of the polymerizing solution to a carbon-coated EM grid, incubate it for 30s, wash it 3 times with miliQ, dryblot and cryoquench in liquid ethane before imaging by 300kV cryoEM. Subsequently obtained micrographs reveal fibril-like particles with a diameter of 5.4±0.4nm (measured from 2D class averages; n=19; standard deviation) and lengths that span from 14nm to hundreds of nm (Fig.4a). Next, we manually picked fibrils shorter than 30nm choosing a box size that fully encloses the raw particles and subjected the extracted particles to multiple rounds of 2D classification in Relion 3.1 (Fig.4b). From this we obtain 2D class averages that correspond to dimeric and trimeric species of R15.5. Due to the sparsity of these particles (4 fibrils on average per micrograph: 1572 picked particles on 362 images), we were not able to clearly resolve any secondary structure elements, however, based on the diameter and the double-ridged features (see^11^ for a protofibril 2D class as reference), we attribute these fibrillar particles to front view projections of an R15.5 dimer and trimer, respectively. We point out that the diameter of the fibrils derived from the class averages (5.4±0.4nm) is larger than the diameter based on the AF2 prediction and high resolution 2D class averages of mature fibrils. We attribute this discrepancy to uncertainties in the particle alignment process during 2D classification as a result of the low number of particles and poor particle contrast, leading to a ‘smeared out’ class average and an overestimated diameter as a result. It is also important to note that we do not observe any other types of particles than amyloid nuclei and fibrils, indicating that R15.5 monomers do not accumulate in any other on- or off-pathway aggregates under these conditions. This is confirmed by the HSQC spectra we collected for R15.5, minutes after buffer exchange (Supporting Figure 8): analogous to CsgA, the R15.5 spectrum is very narrow in the proton dimension (8.7-7.8ppm) in the absence or presence of 10mM CaCl_2_, indicative of the unstructured nature of the R15.5 monomer in its pre-amyloid state.

### Canonical curlin strands are enriched towards amino acid types that have low **β**-strand propensity

Multiple sequence alignment (MSA) analysis of curlin repeats has uncovered strong co-evolutionary couplings that have led to an unambiguous prediction of the curlin fold, i.e. a β-solenoid^28^. That prediction has since been validated for CsgA by solid-state NMR^29^ and more recently via cryoEM^11^. And yet, despite the all-β nature in the finally folded state, CsgA^19^ and CsgA^slowgo^ (this work) both exhibit a predominant helical character in their respective pre-amyloid forms – an unexpected secondary structure propensity that was not predicted by co-evolutionary couplings analysis^28^ and subsequent neural-network based models (AlphaFold2, RosseTTAFold) that leverage this information that is embedded in the MSAs for structural prediction purposes (e.g. entry A0A5C9ANS0 in the AlphaFold Protein Structure Database and ^30^). To understand this observation, we performed a sequence analysis of the curlin motif focusing on the relative abundances of the amino acids in said curlin sequences and cross-referenced those to amino acid secondary structure propensities derived from PDB analyses. In Figure 5, we show a stacked column plot of the residue types found in positions 1 to 8 of the motif A and B β-strands of the curlin repeats of CsgA and R15.5^31^, averaged over all repeats, using a colour scale that reflects the mean amino acid propensity for β-strand conformations as defined in Fujiwara *et al*^32^. Positions 1, 3, 5 and 7 map onto inwards facing residues of the strand and follow the canonical curlin strand motif N-$-Ψ1-$-Ψ2-$-Q where Ψn are hydrophobic residues (A,I,V,L,F), S or T. As we^11^ and others^33^ have previously shown that the steric zipper (inward-facing) residues of the curlin fold are well conserved, we focus here on the outwards-facing residues of the strands of the β-solenoid. We find that positions 2, 4, 6 and 8 of CsgA and R15.5 are predominantly populated by amino acid types that have comparatively low β-strand propensities. To quantify this, we calculated the weighted average of the β-strand propensity ∑ f_i_ P_i_ at each position along the curlin strands, with *f_i_* the relative abundance of amino acid of type *i* and *P_i_* the corresponding β-strand propensity for that amino acid type. Fujiwara *et al* refer to *P*_β_ > 1.1 as a *β-strand-favored* amino acid type, *P*_β_ < 0.9 as a *β-strand breaker* and 0.9 < *P*_β_ < 1.1 as neutral. Next, we averaged this across all strand positions to determine the mean β-strand propensity <*P*_β_> for the sequences of CsgA and R15.5 (not factoring in the loop regions), yielding <*P*_β_>_CsgA_= 1.00 and <*P*_β_>_R15.5_= 0.99. To expand the analysis beyond CsgA and R15.5, we mined the RefSeq database for CsgA-like repeat sequences using the hidden Markov model for the curlin repeat from Dueholm *et al*^33^. Based on a curated MSA of 53574 thusly obtained repeat sequences, we constructed the *curli consensus* sequence shown in Fig.5. Similarly to CsgA and R15.5, we find that a canonical curli strand follows a similar secondary structure propensity pattern along the consecutive positions of the strand, and that it is not exclusively composed of high β-strand propensity residues, but strikes a neutral balance with <*P*_β_>_Consensus_ = 1.06. To place this in perspective, we looked for β-solenoidal proteins that have no polymerizing or amyloidogenic traits. For instance, the ice binding protein of *Lolium perenne* shares remarkable structural similarities with CsgA (Supporting Figure 9) but has no reported propensity to form fibrillar structures. Interestingly, we obtain a value of 1.27 for <*P*_β_>_3ULT_, demonstrating relative enrichment of β-strand prone residue types.

## Discussion

Our combined NMR, nMS and cryoEM data for CsgA, CsgA^slowgo^ and R15.5 shed further light onto the nucleation mechanism of the functional amyloids found in the extracellular matrix of Gram-negative prokaryotes. We find that the major curlin subunit is only partially unfolded in its monomeric state, residing in a condensed state that is sparsely sampling local secondary structure. This agrees with NMR data published by Sewell and coworkers on the pre-amyloid ensemble of CsgA^19^. Notably, we found no indications that CsgA or R15.5 exist as folded monomers, demonstrating that folding is tightly coupled to polymerization. This leaves open two possible scenarios (Fig.6). In the first scenario, the rate-limiting step in fibril formation is the folding of CsgA into its β-helical structure which then serves as a catalytic center for newly incoming unfolded CsgA monomers to dock and fold cooperatively. Consequently, folded CsgA monomers would be rare and short-lived species and the system will have a nucleus of size 1. The second scenario that is compatible with our observations would be one where only a folded dimeric protofibril (nucleus of size 2) is thermodynamically stable, requiring monomers to fold and dock either cooperatively or in short succession. Andreasen and coworkers^17^ obtain a value of 1.12 for the reaction order of the primary nucleation process, suggesting that scenario 1 is more likely for CsgA.

**Figure 6:**
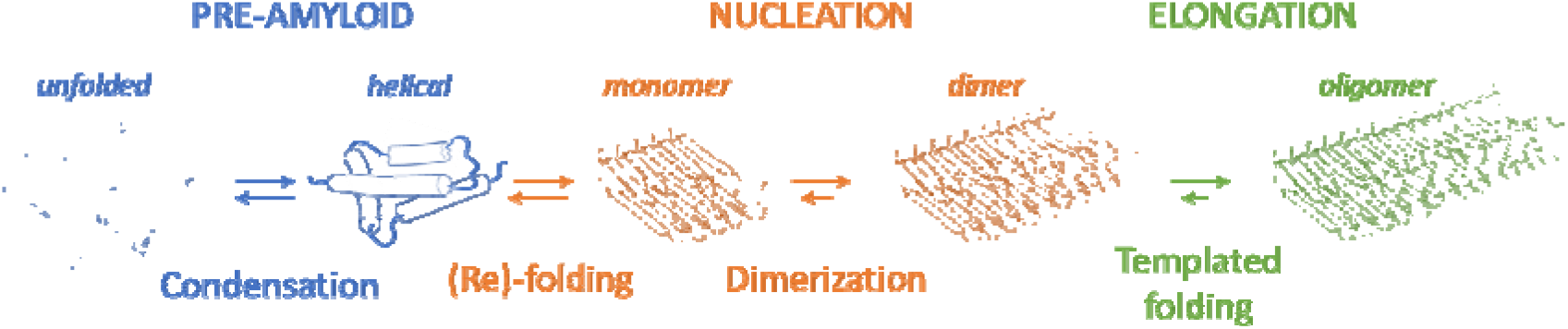
Model for the formation of curlin fibrils starting from an unfolded monomer that first condenses into a molten globule-like state with predominant helical character. The slow refolding of this pre-amyloid CsgA into a β-solenoidal monomer forms the primary stage of nucleation. Next, two Csga monomers join to form a dimeric nucleus, which constitutes a minimalistic curlin protofibril. The termini of this protofibril then serve as catalytic surfaces for the templated folding of incoming subunits during the stages of fibril elongation.

The difference between a monomeric or dimeric curlin nucleus is perhaps of secondary importance, precisely because the nucleation mechanism may change depending on the location in the phase diagram. For instance, Morel and coworkers^5^ have argued that Ab peptides can switch from a nucleated conformational conversion process when the peptide concentration is above the critical micelle concentration (CMC) to a bimolecular nucleation process characterized by linear nuclei below the CMC. The reaction order of primary nucleation could be a function of both external (i.e. solutal composition and reaction conditions) and/or intrinsic factors such as the number of repeats in the primary sequence. For instance, we detect 60 curlin repeats in the primary sequence of the hypothetical protein FJ977_01705 (accession: TPN01251.1) found in the genome of *Mesorhizobium* sp. B2-1-3A, and locally integrated into a CsgG-A-E-B-F operon. It is perhaps likely that the folding energy associated with the formation of a 60-stranded β-solenoid would be sufficient to stabilize the folded monomeric state. More relevant is the conceptual difference with respect to the nucleation pathways of pathological peptide amyloids that involves inter-molecular bonding into pre-amyloid aggregates from which the amyloid structure needs to be ‘found’ by sampling phase space. For CsgA that search can be viewed as a folding problem, where folding essentially equates to nucleation. Moreover, there is no reported polymorphism for CsgA amyloids despite more than a decade of active research on it and we have recently shown that CsgA and R15.5 fibrils can be considered isomorphous^31^, sharing the same steric zipper mechanism to drive the folding reaction as suggested by the high degree of conservation of the corresponding residues. This consistency in protofibril structure over a wide range of conditions is compatible with CsgA’s ‘nucleation-via-folding’ mechanism, i.e. the fibril structure is encoded in the primary sequence of the subunits, and in particular the conserved, inward facing residues that make up the steric zipper. This stands in sharp contrast to the structural variety that can be found for pathological amyloids composed of otherwise identical proteins^34^.

For curli, structural consistency facilitates promiscuous cross-seeding between curli produced in interspecies biofilms^35^. In addition to structural reproducibility, the folding-limited kinetics is likely also a result of biological selective pressures. Produced in the cytoplasm, CsgA needs to be transported across the inner membrane, traverse the periplasm, and be shuttled through the outer membrane pore CsgG, all while remaining in its pre-amyloid state. Although the cell has coping mechanisms in the form of two chaperones (Cytosol: DnaK^36^; Periplasm: CsgE^37^) and one inhibitor (CsgC^38^ or CsgH^39^) to prevent and block untimely polymerization, it has been shown that finetuning of the rate of CsgA folding yields additional spatiotemporal control over curli formation to avoid cytotoxic effects^40^.

Indeed, the general concept of kinetic control for CsgA folding was introduced by Wang *et al*^40^ in the form of gatekeeper residues^41^ and more recently in^26^. Although Wang *et al* did not propose a molecular underpinning for this process, NMR observations of the pre-amyloid state of CsgA now allow us to formulate a hypothesis for this mechanism. Specifically, the predominant helical character of the pre-fibrillar CsgA monomers points towards the rapid emergence of local regions of helical secondary structure when CsgA transitions from denaturing to native buffer conditions (Fig.6) – a folding pathway that was unexpected given the all-β character of the folded state. Our sequence analysis shows that the CsgA sequence and its homologues strike a balance between residues with high β-strand propensity (V,I,T,Y,F) and residues that tend to be underrepresented in strand regions (G,A,P,D,N,Q). Given that the amyloid curlin kernel **N**/S-X-**A**/S-X-V/I-**Q**-X-**G** largely consists of low β-propensity residues, one could naively expect that the net ‘β-balance’ would be restored via the amino acid distribution at the X-residue sites. Considering that these sites are surface exposed and therefore not controlled by the structural restraints of the curlin core, one might assume that the selective pressures are weaker for these sites. Our sequence analysis shows that this may not be the case, rather positions 2, 4, 6 and 8 in Fig.5 are not enriched towards β-favoured residues. Taken altogether, a picture emerges of a folding landscape with a local minimum that corresponds to a molten globule-like state with helical character. The local restructuring from helix to strand could be an activated step that represents a bottleneck (rate-limiting) step on the path towards the folded β-solenoidal state.

## Materials and Methods

### Protein production and purification

CsgA (uniprot: P28307), CsgA^Q49A/N54A/Q139A/N144A^ (i.e. CsgA^slowgo^) and R15.5*.5* (uniprot: A0A0E3UX01) were cloned into pET22b via the NdeI site without their signal sequence but with a C-terminal tag 6xHis-tag. Expression was induced in BL21 *slyD^-^* or C43(DE3) cells by addition of 1 mM IPTG at an OD_600_ of 0.6. Cells were harvested by centrifugation at 5,000 g for 10 minutes after one hour of induction. Cell pellets were lysed for 30min in buffer A (50 mM Kpi pH 7.2, 500mM NaCl, 8 M urea, 12.5mM imidazole) and the cell lysate was centrifuged at 40,000 g for 30 minutes at 20°C. After sonication on ice (Sonics Vibracell sonicator: pulse 30s/30s on/off; 50% amplitude; 3min total time; VC130PB 50-100mL solid probe) to reduce the viscosity of the lysate, the supernatant was loaded on a HisTrapTM FF column (GE Healthcare Life Sciences) equilibrated with 5 column volumes (CV) of buffer A. After washing in 10 CV buffer A, the protein was eluted using buffer B (50 mM Kpi pH 7.2, 8 M urea, 250 mM imidazole) and fractions were analyzed via SDS-PAGE. CsgA preparations that were purified from C43(DE3) contained SlyD as a minor contaminant. Relevant protein fractions were pooled and filtered with a 0.22µm cutoff filter to remove any potential amyloid seeds and stored at –80°C.

To prevent any unwanted amyloid formation in our protein stock solutions, all purification steps were performed under denaturing conditions (8 M urea) and handling times at room temperature were reduced to an absolute minimum. This approach allows us to store CsgA and R15.5 in their pre-amyloid, unfolded form and gives control over the exact starting point of polymerization by buffer switching to native conditions, i.e. 15 mM MES pH 6.0. To remove urea, ZebaTM Spin Desalting columns (7K MWCO) (Thermo Scientific) or 5mL HiTrap Desalting Columns (GE Healthcare) were used.

### Thioflavin T assays

ThT assays were performed using freshly desalted CsgA in 15 mM MES pH 6.0. Protein samples were pipetted into a black flat-bottom 96-well microplate (Greiner Bio-One) in the presence of 50 μM Thioflavin T dye (ThT). Fluorescence measurements were performed using the Infinite 200 plate reader (Tecan) at 25 °C with excitation at 430 nm and emission at 495 nm. Fluorescence readings were taken every 10 min, and the plate was shaken for 5 s before each reading.

### NMR

Unless otherwise stated, all NMR experiments were carried out at 25 °C. Samples of CsgA^slowgo^ were produced with uniform 15N or 15N/13C labelling. This was carried out by cell growth in LB until the cells had reached and OD_600_ of 0.6 before centrifugation and resuspension of the pellet in M9 growth media containing ^15^N ammonium chloride and ^13^C-glucose before induction of expression. For the time course experiments, unlabelled CsgA, CsgB and CsgA^slowgo^ samples in 8 M urea buffer were taken and run over a Hitrap Desalting column into 15 mM MES buffer at pH 6.0. The samples were then transferred immediately into a 5 mm NMR tube (Wilmad) and placed into a 600 MHz magnet with a coldprobe (Agilent). The peptide backbone was assigned using a combination of BEST versions of 3D HNCA, HNCOCACB, and HNCO spectra^42^ and ^15^N NOESY, ^15^N TOCSY, HNCA, CBCACONH, HNCO, HN(CACO)NH and H(NCACO)NH experiments on a 300 μL sample of 100 μM 13C/15N CsgA^slowgo^ in 15 mM MES pH 6.0 buffer in a 5mm Shigemi sample tube (Shigemi ltd). T1, T2, heteronuclear NOE, CLEANEX and DOSY data were collected on 100 μM ^15^N-labelled protein at a proton Larmour frequency of 600 (Agilent) or 800 (Bruker) MHz. Data were processed using NMRpipe^43^ and analysed using CCPN Analysis^44^. Data were fit using peakypy^45^ and nmrglue^46^ and in-house Python and Nmrpipe scripts. Predicted shifts for the structural models were calculated from PDB data calculated from the sequences of CsgA and R15.5 by Alphafold 2 using the ShiftX2 server^47^. The 1D ^1^H spectra of R15.5 were recorded every 10 min for the total duration of 10 hours at 298 K on a Bruker Avance III HD 800 MHz spectrometer, equipped with a TCI cryoprobe. The sample contained 6% D_2_O for the lock. The NMR data were acquired, processed and analysed in TopSpin 3.6 (Bruker).

### Native mass spectrometry

Native MS experiments were performed on a Synapt G2 High Definition mass spectrometer (Waters Corporation, Wilmslow, UK) with travelling wave (T-wave) ion mobility and equipped with a nano electrospray ionization (nESI) source, using *in-house* generated gold-coated borosilicate capillaries. The critical experimental parameters were set at: capillary voltage 1.4–1.6 kV; source temperature 30 °C; sample cone 50 V; extraction cone 10 V; transfer collision energy 10 V; trap bias 45 V; IM wave velocity 300 m/s; IM wave height 35 V. Native mass spectra were acquired at trap collision energy of 45 to 60 V. For collision unfolding experiments, measurements were made at trap accelerating voltage steps of 10V from 5V to 100V. Gas pressures in the instrument were: backing 8 mbar, trap cell 0.04 mbar; IM cell 2.1 mbar; transfer cell 0.036 mbar. CsgA and CsgA^slowgo^ samples were buffer-exchanged into MS-compatible buffer (150 mM Ethylene diamine diacetate pH 6.3, 10mM TMAO) using Micro Biospin 6 desalting columns (Bio-Rad) to a final protein concentration of 3-5 μM. The individual proteins samples were immediately loaded into gold-coated nanoflow capillaries and introduced into the mass spectrometer by nESI. Mass measurements were calibrated using cesium iodide (100 mg/mL). Spectra were recorded and smoothed using Masslynx 4.1 (Waters) software. Gas-phase collision induced unfolding analysis was performed using the software PULSAR^48^.

### Cryo-EM grid preparation and image acquisition

High-resolution cryo-EM datasets were collected using continuous carbon coated Quantifoil™ R2/1 400 copper mesh holey carbon grids. Grids were glow-discharged at 5 mA plasma current for 1 minute in an ELMO (Agar Scientific) glow discharger. Gatan CP3 cryo-plunger set at room temperature and relative humidity of 90 percent was used to prepare the cryo-samples. 3 µL of a polymerizing R15.5 solution was applied on the CC grid and incubated for 30 seconds. The solution was dry-blotted from both sides using Whatman type 2 paper for 3 seconds with a blot force of 0 and plunge-frozen into precooled liquid ethane at -176 °C. High resolution cryo-EM 2D micrograph movies were recorded at 300 kV on a JEOL Cryoarm300 microscope equipped with an in-column Ω energy filter (operated at slit width of 20 eV) automated with SerialEM 3.0.8^49^. The movies were captured with a K3 direct electron detector run in counting mode at a magnification of 60k with a calibrated pixel size of 0.764 Å/pix, and exposure of 64.66 e/Å^2^ taken over 61 frames. In total 362 movies were collected within a defocus range of 0.5 to 3.5 micrometers.

## Author contributions

MS and HR designed the project. BP performed cryogenic freezing, cryoEM imaging and data processing. CM collected nMS data and performed the analysis with input from FS. JKC collected all NMR data and performed the analysis. MS wrote the manuscript with contributions from all authors.

## Competing Interests Statement

The authors declare to have no competing interest.

## Data Availability Statement

The data related to this manuscript is available upon request to MS and/or HR.

## Acknowledgements

We thank Marcus Fislage and Dirk Reiter at the VIB-VUB Facility for Bio Electron Cryogenic Microscopy (BECM). This work was funded by VIB, EOS Excellence in Research Program by FWO through grant G0G0818N to HR and G043021N to MS.

---

**Supporting Figure 1:**
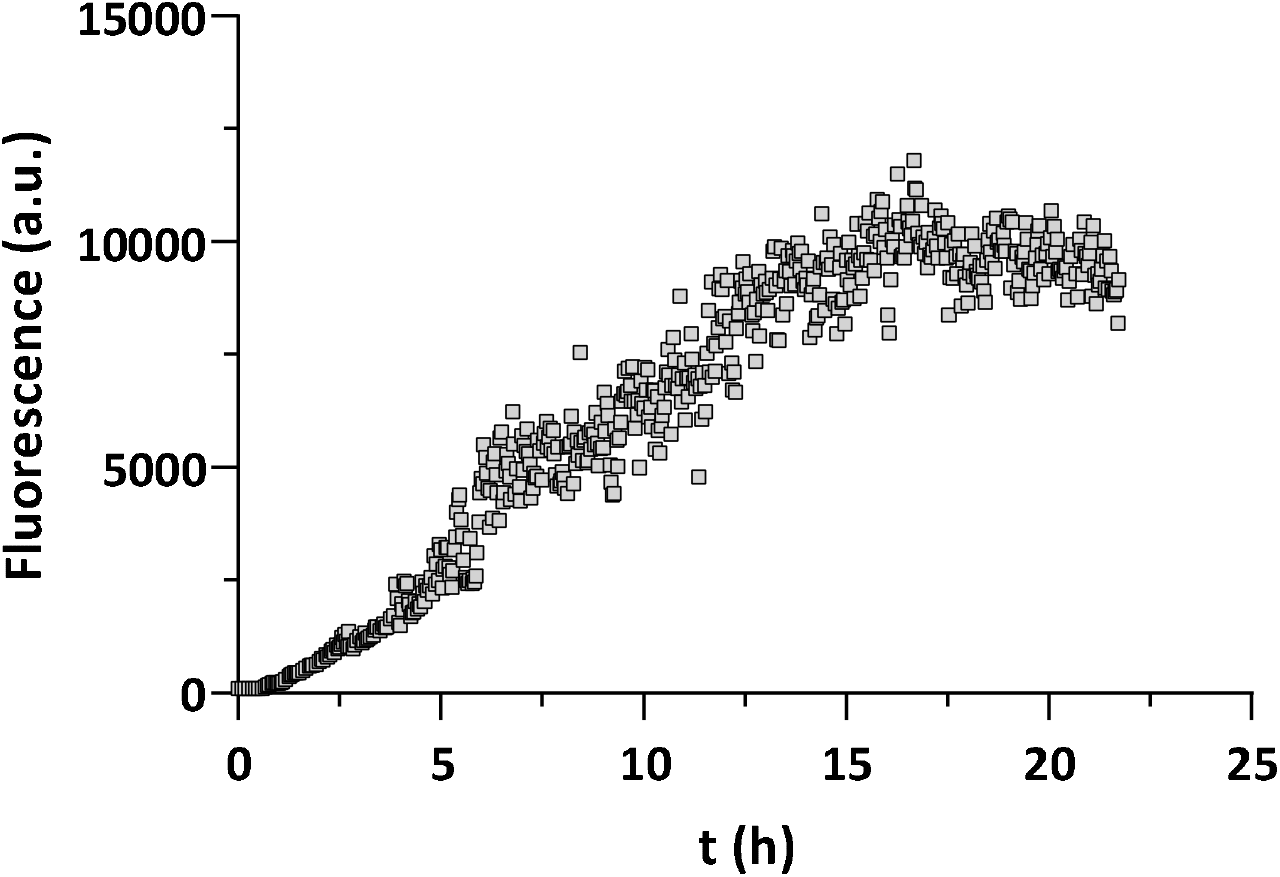
Tht-fluorescence of a freshly desalted 5µM CsgA solution in 15mM MES pH 6.0 at 20°C.

**Supporting Figure 2:**
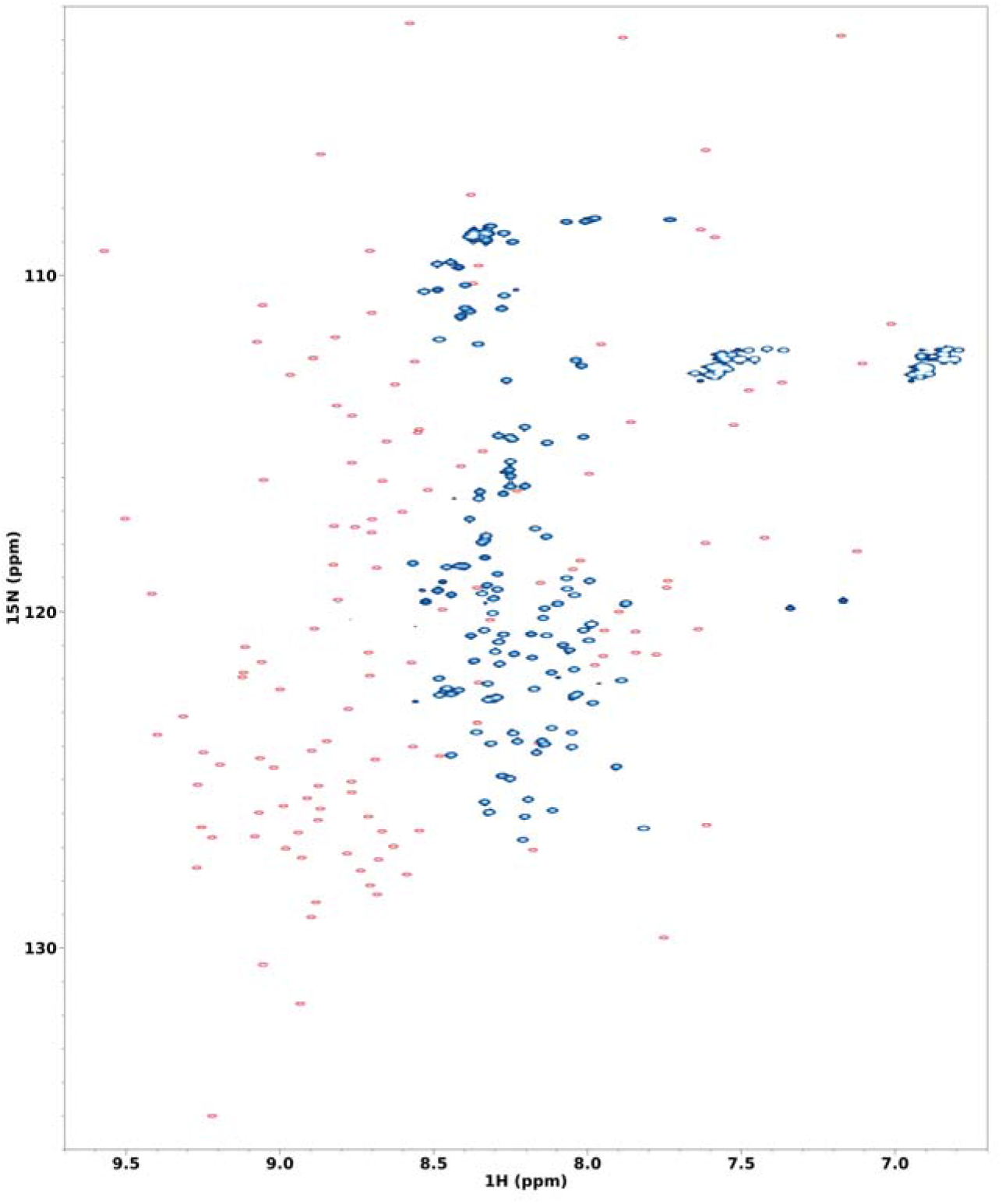
Overlayed ^15^N,^1^H BEST-HSQC spectrum of CsgA^slowgo^ (blue peaks) compared with the shifts calculated from the predicted structure (red peaks). This shows the low dispersion of the acquired spectrum in comparison to that of the fully folded CsgA spectrum in the prediction.

**Supporting Figure 3:**
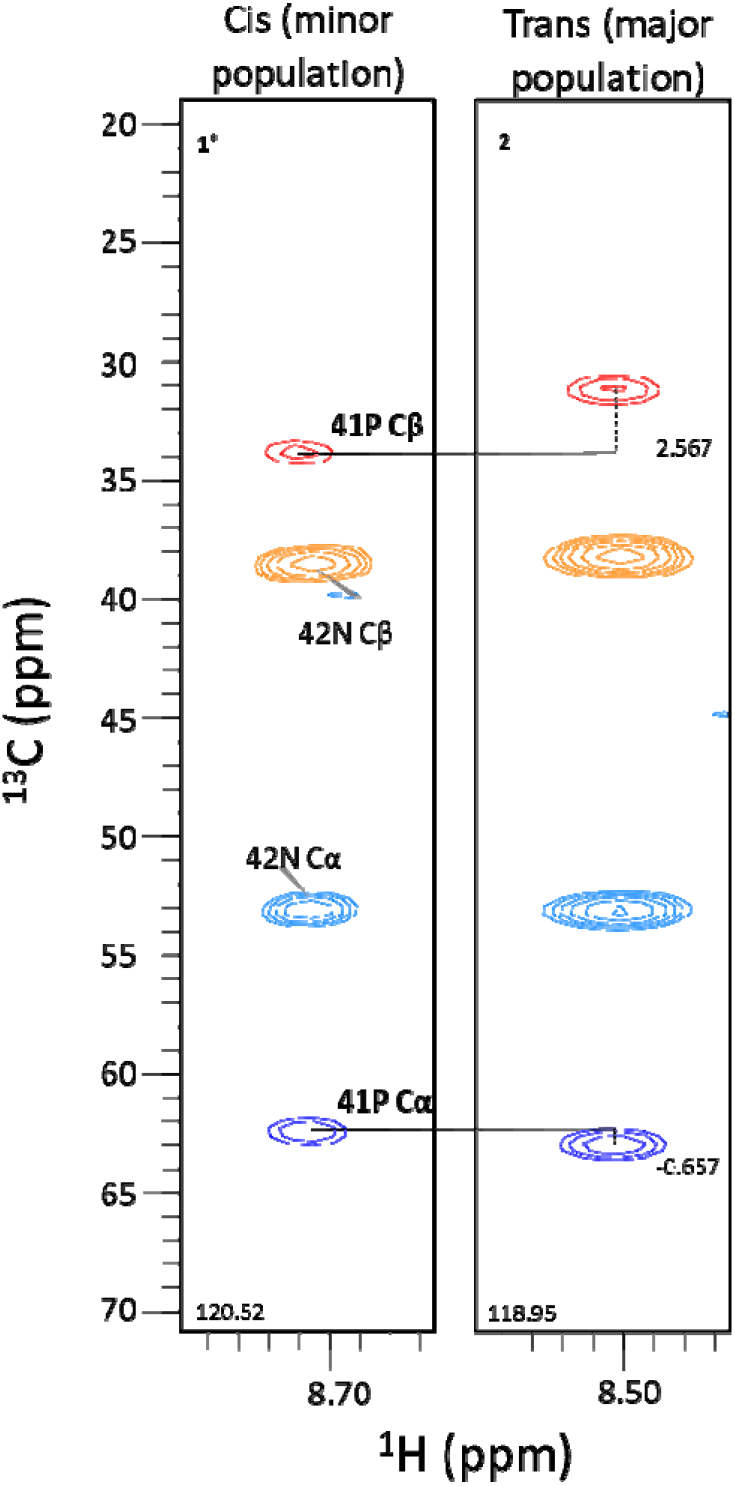
Strips from CBCA(CO)NH spectrum of CsgA^slowgo^ for 42N (right) and 42N^minor^ (left) showing the CC and Cβ peaks for 42N (i) and the (i-1) residue, 41P. Proline side chain resonances are sensitive reporters on the proline *cis-trans* isomerisation state. The downfield shift of the Cβ peaks and the upfield shift of the CC peaks for 41P^minor^ is indicative of the *cis* configuration.

**Supporting Figure 4:**
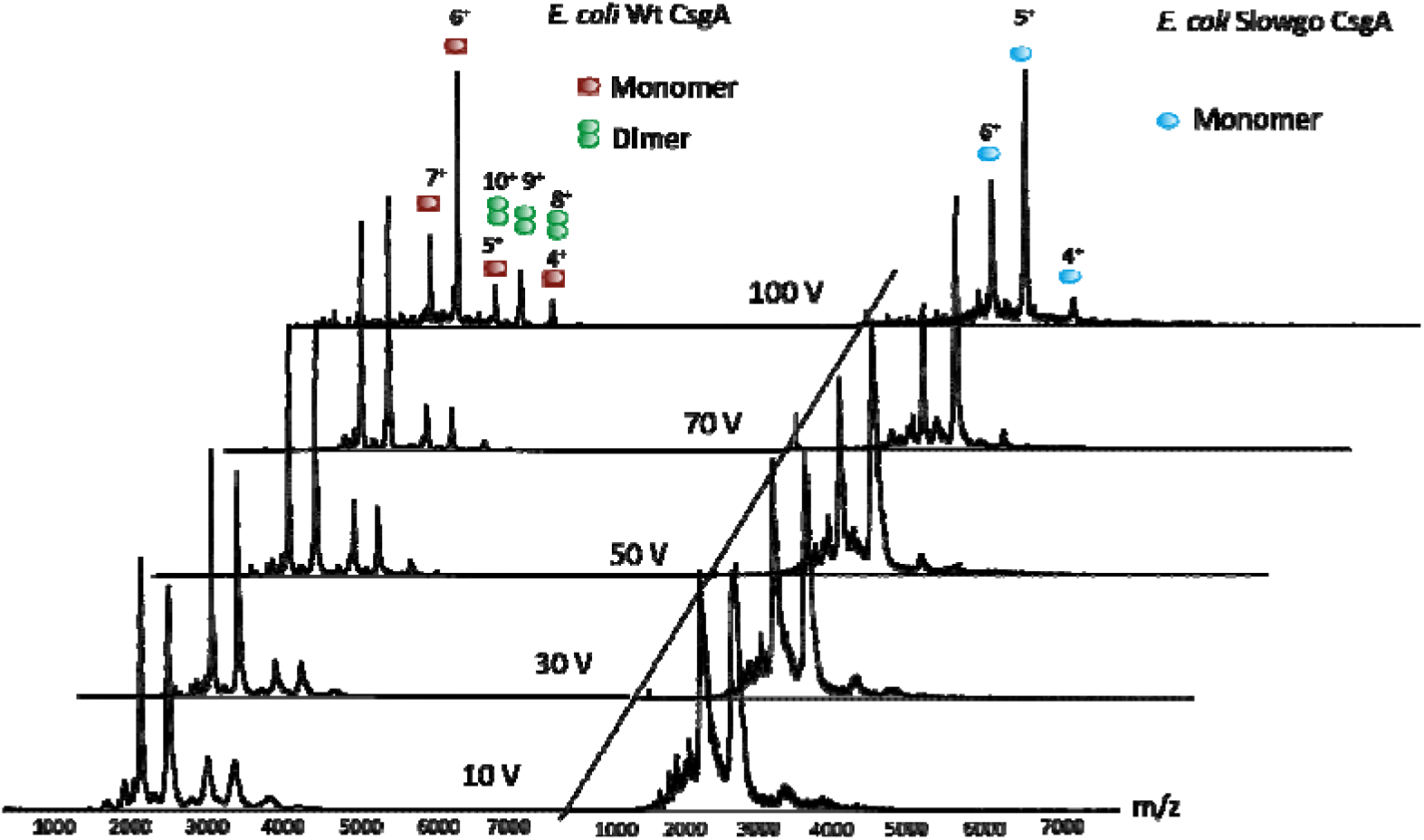
Mass spectra of WT CsgA (left panel) and CsgA^slowgo^ (right panel) under native conditions at increasing trap collision voltage values. At 100V, the different peaks are well resolved and allow unambiguous assignment to monomeric (red and blue) and dimeric CsgA (green) species respectively.

**Supporting Figure 5:**
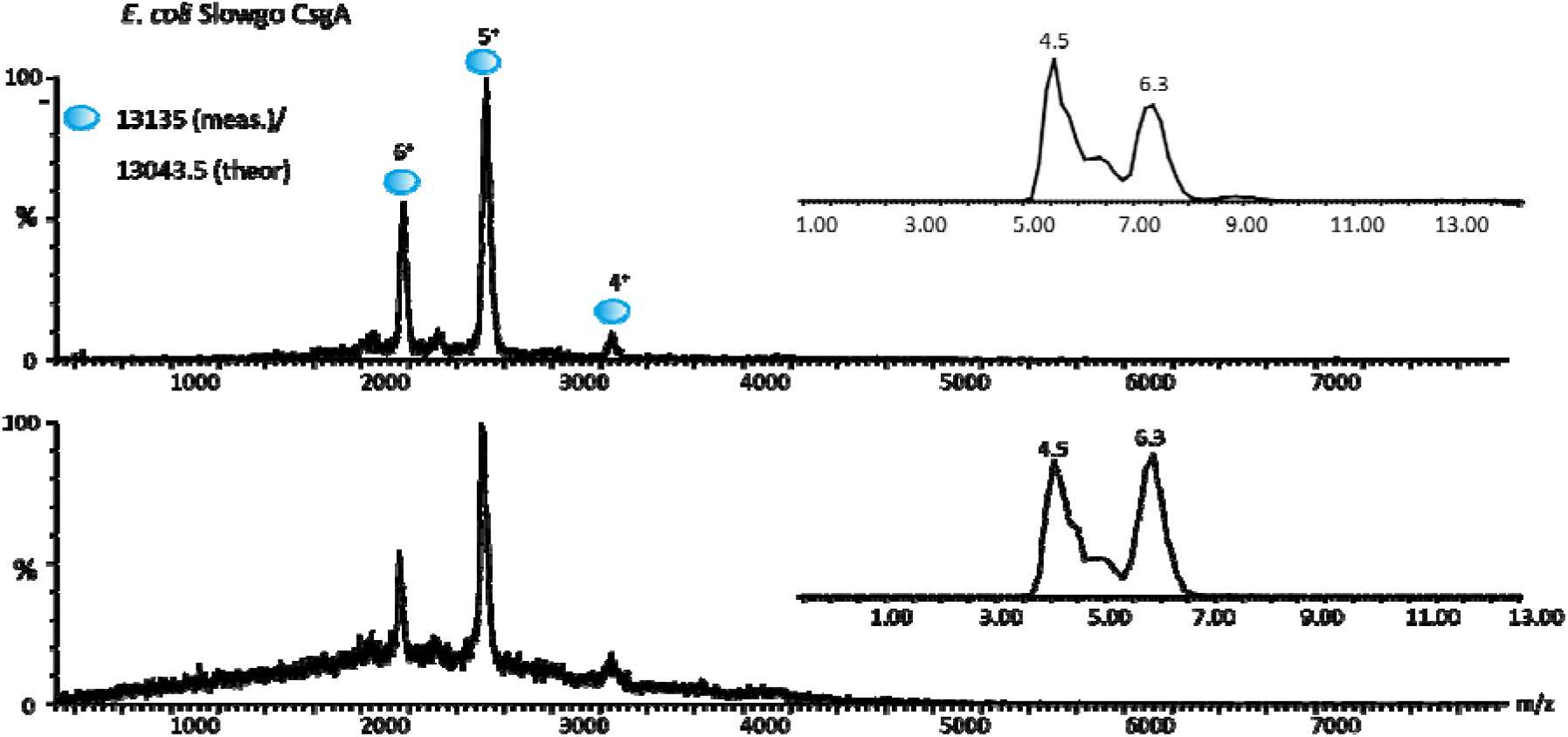
Mass spectra of CsgA^slowgo^ immediately after buffer exchange (top panel) or after 2h on ice (bottom panel). The inset shows the drift time of the 6+ charge state peak. Two species with different gas-phase mobilities are present in the sample, right after the exchange but also after the 2h waiting time, indicating no major conformational change / degradation of the sample over time.

**Supporting Figure 6:**
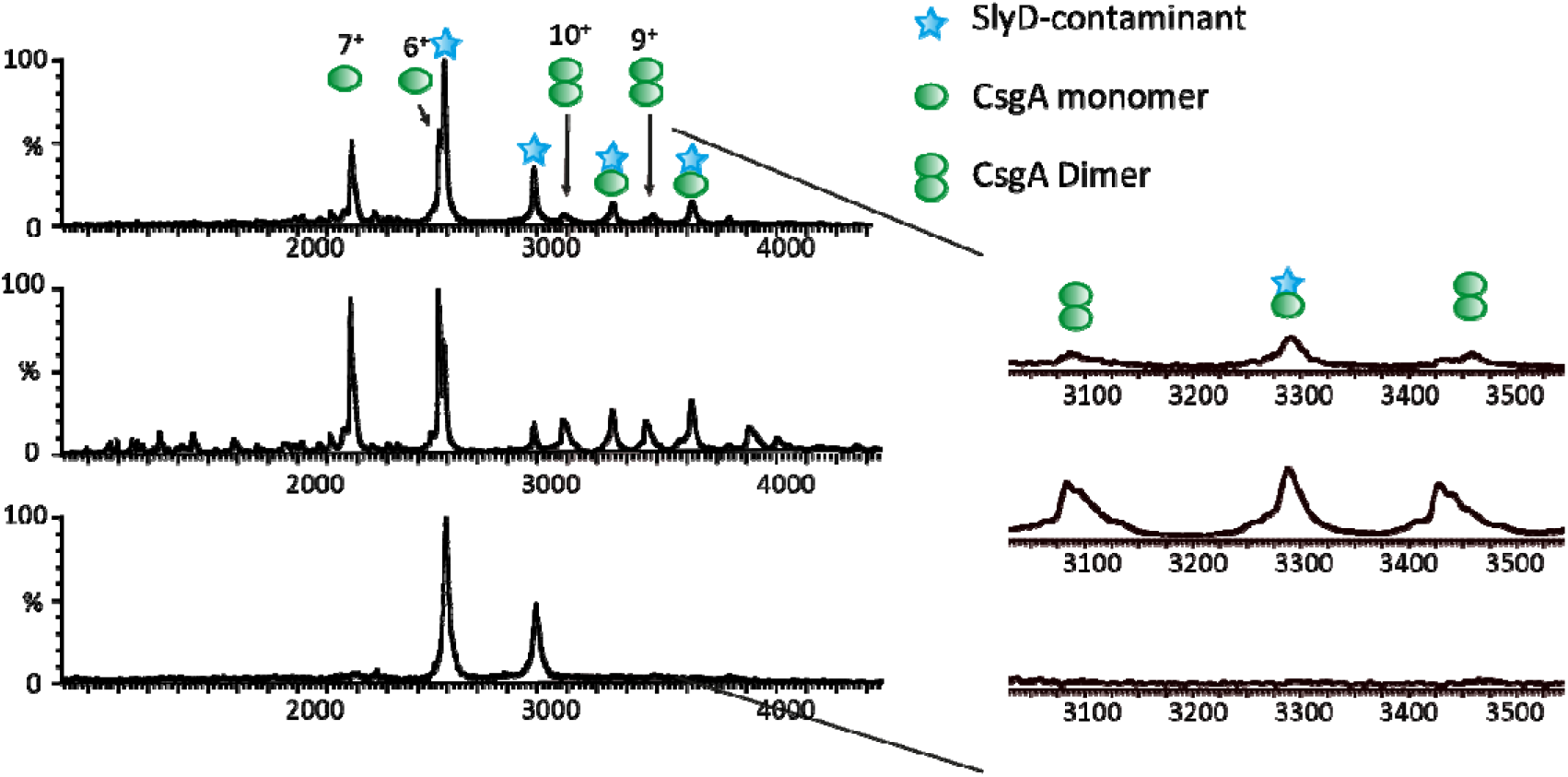
Mass spectra of WT CsgA directly after buffer exchange (top panel), after 40 min on ice (middle panel) and after 1h on ice (bottom panel). After 1hour, only the peaks corresponding to the contaminant SlyD can be detected. The inset on the right shows that the intensity of the dimer peak increases from 0 to 40 min, then can no longer be detected.

**Supporting Figure 7:**
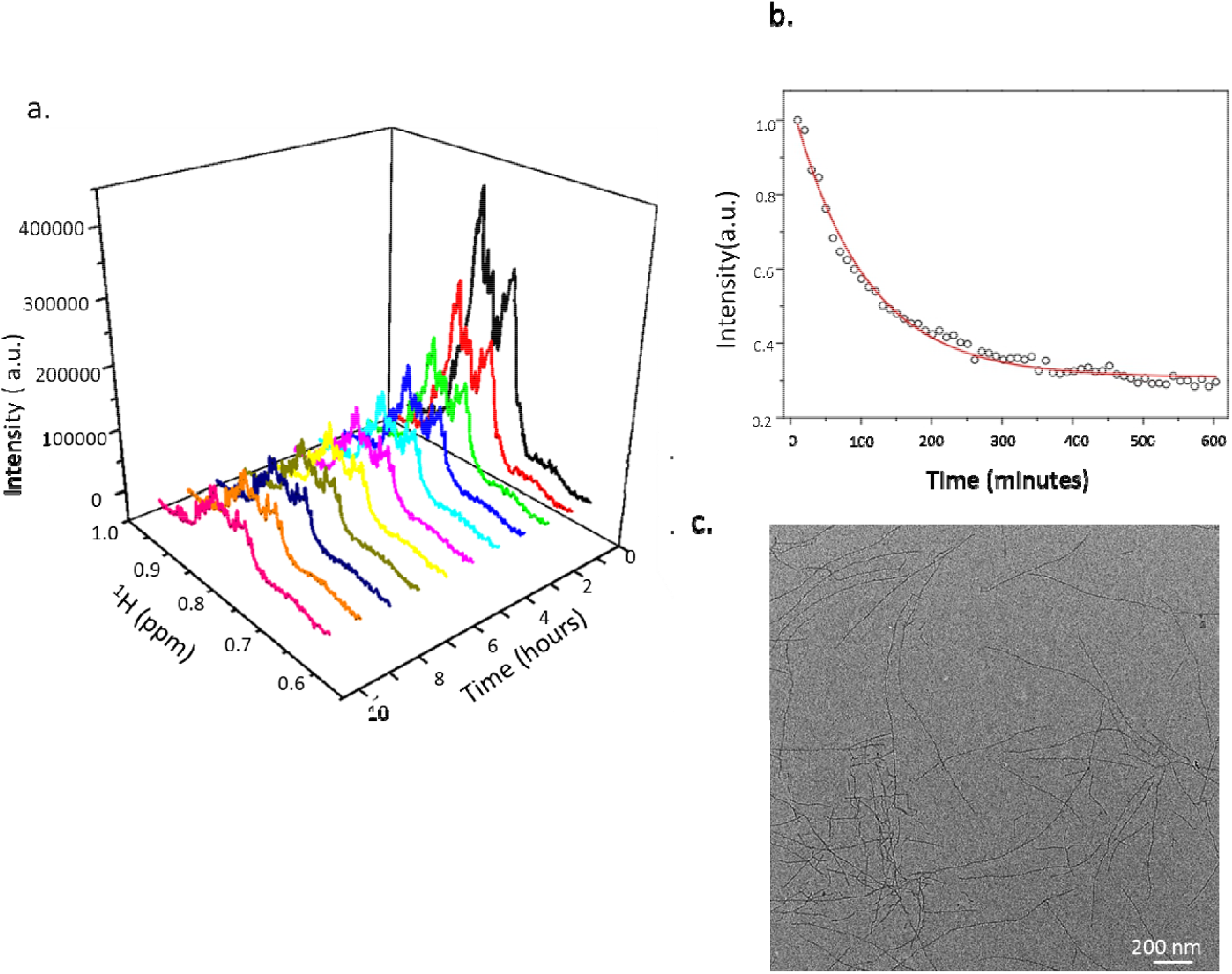
**(a)** Time-dependent spectral changes in the methyl region of 1D ^1^H NMR spectrum of R15.5 after buffer exchange to citrate-phosphate buffer at pH 7.0. **(b)** Normalized NMR signal intensity, defined as the integral over the spectral region displayed in (a), as a function of time. The red curve is the best fit to a single exponential decay function with a decay rate of 0.58 ± 0.02 h^-1^. (c) Negative stain micrograph of R15.5 taken 12 hours after buffer exchange to non-denaturing conditions.

**Supporting Figure 8:**
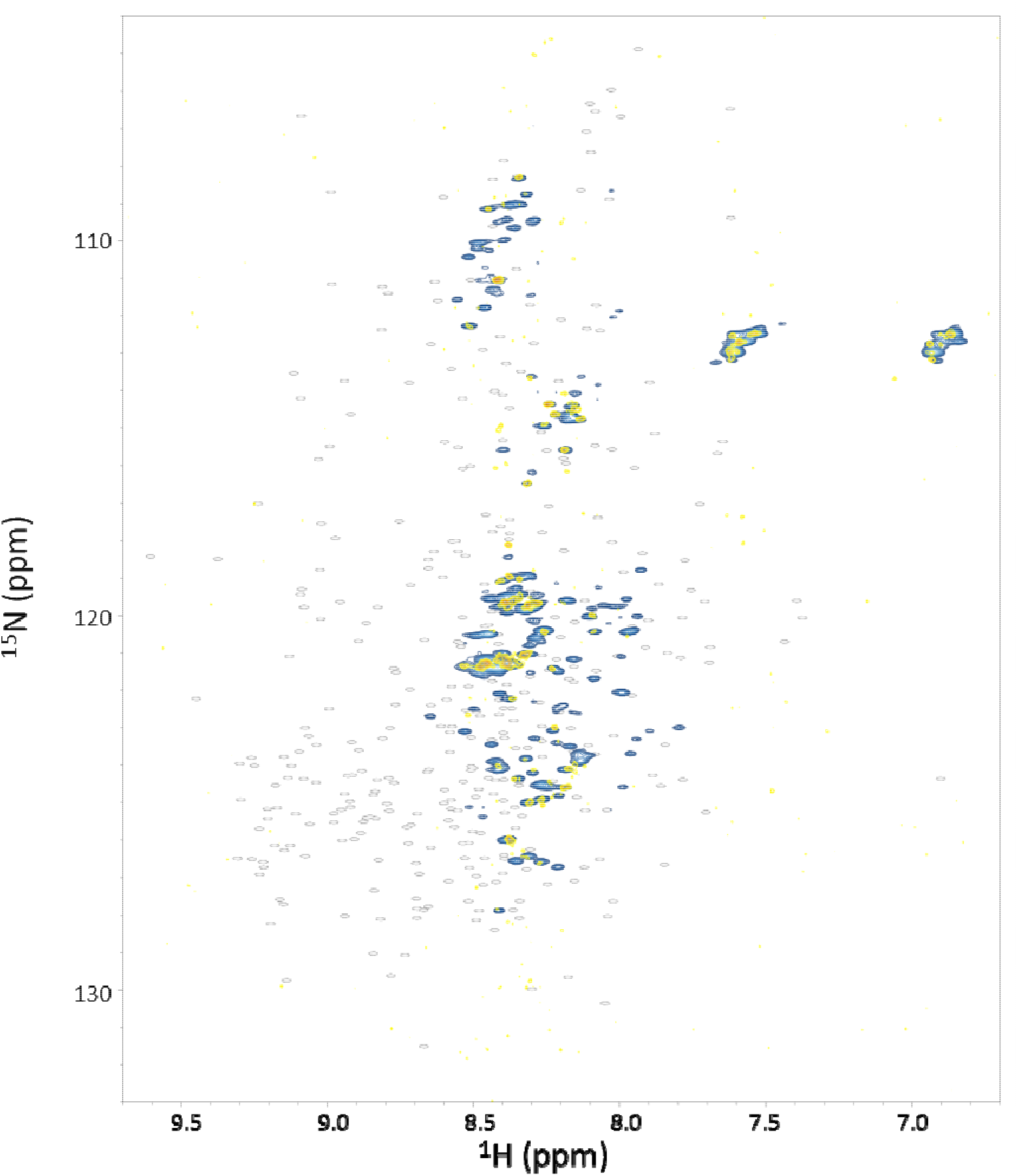
Overlayed ^15^N,^1^H BEST-HSQC spectra of 100 µM ^15^N-labelled R17 in 15 mM MES buffer at pH 6.0 (blue) and in 15 mM MES buffer at pH 6.0 with 10 mM CaCl_2_ (yellow) to probe for divalent cation mediated folding. The narrow dispersion of the amide peaks in the proton dimension is indicative of disorder. There is significantly less signal observable in the spectrum acquired in the presence of CaCl_2_. This is likely to be due to the rapid formation of amyloid structures and subsequently larger species that are now too large to be observable in solution-state NMR. Predicted shifts for the folded state of the protein are shown as grey ellipses.

**Supporting Figure 9:**
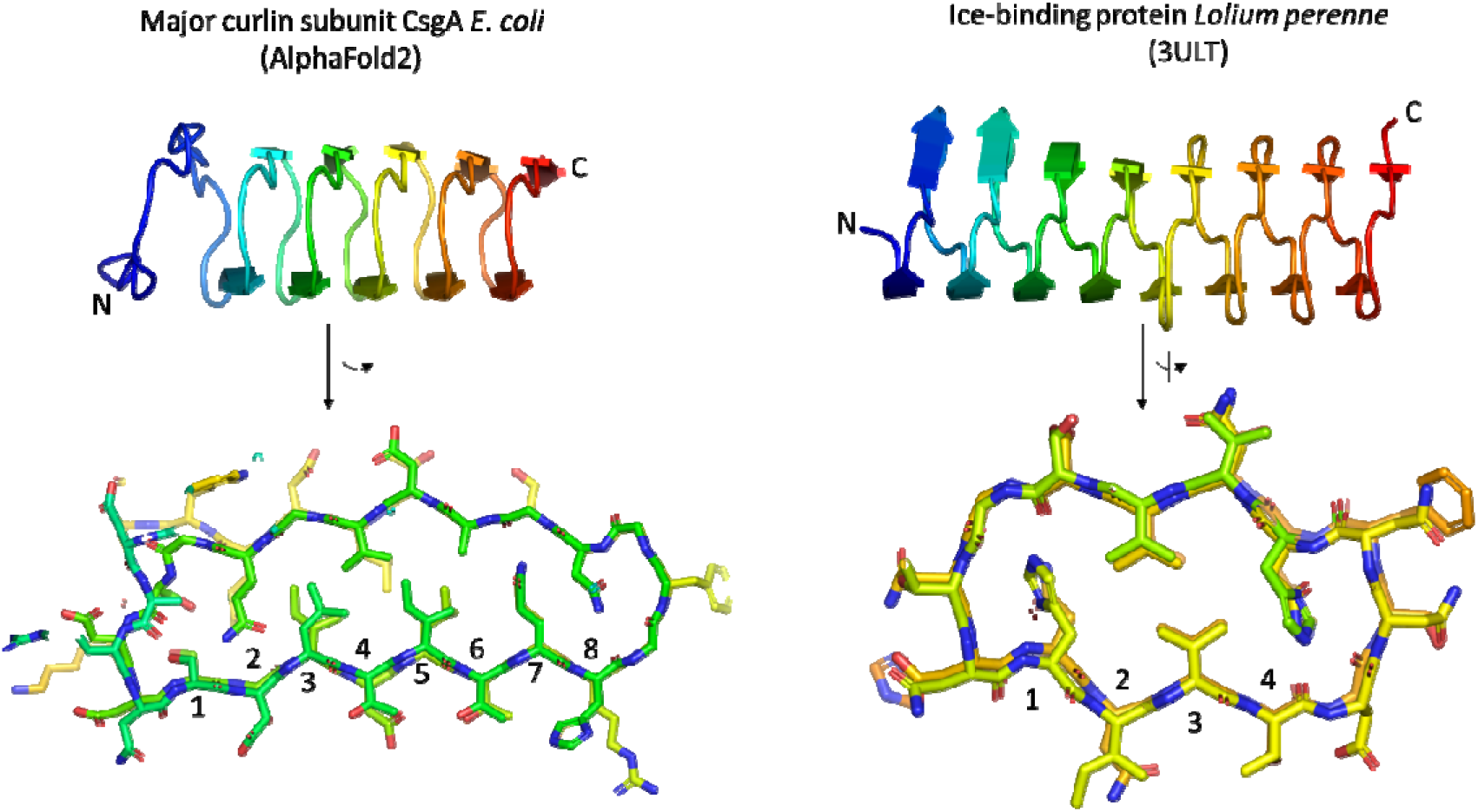
Cartoon side-view and cross-sectional view in stick representation of CsgA *E. coli* and the β-solenoidal ice-binding protein from *Lolium perenne*. Numerical labelling of the positions along the β-strand correspond to the nomenclature used in Figure 5.

## Notes

### Competing Interest Statement

The authors have declared no competing interest.

